# Departure from OFF-State Microstate Dynamics Tracks Levodopa Response in Parkinson’s Disease

**DOI:** 10.64898/2026.08.03.742400

**Authors:** Matteo Demuru, Marianna Angiolelli, Emahnuel Troisi Lopez, Mario De Luca, Enrica Gallo, Domenico Tafuri, Damien Depannemaecker, Carmine Granata, Giuseppe Sorrentino, Pierpaolo Sorrentino

**Affiliations:** Department of Medical, Motor, and Wellness Sciences, University of Naples “Parthenope”, Naples, Italy; Department of Education and Sport Sciences, Pegaso University, Naples, Italy; Institute of Systems Neuroscience, Aix-Marseille University, INSERM, UMR1106, Marseille, France; Institute of Applied Sciences and Intelligent Systems, CNR, Pozzuoli, Italy

## Abstract

Although dopaminergic therapies in Parkinson’s disease primarily restore dopamine within nigrostriatal circuits, symptoms are better indexed by whole-brain dynamics than by local activity. In this manuscript, we hypothesize that L-Dopa therapy affects whole-brain dynamics alike, which in turn relate to clinical improvement. To test this hypothesis, we conducted a repeated-measures, source-reconstructed MEG study in 13 bradykinetic-dominant PD patients, recording resting-state cortical activity OFF medication and ∼1 hour after levodopa administration (ON). We characterize brain dynamics using a microstate framework, in which transition probabilities between microstates are used to contrast pathological OFF-state dynamics with those in the ON-state. Microstate dynamics were stable within medication states but reconfigured by L-Dopa, with greater departures from the OFF-state pattern associated, at the individual level, with larger clinical improvements. Our results suggest that individualized changes in microstate dynamics may serve as a neurophysiological marker of dopaminergic responsiveness.

## Introduction

Parkinson’s disease (PD) is a multi-system neurodegenerative disorder characterized by a heterogeneous clinical profile encompassing both motor and a broad spectrum of non-motor symptoms ^1,21,2^. Depletion of dopaminergic neurons in the pars compacta of the substantia nigra is a key hallmark of the disease, and pharmacological therapy aims to supplement dopamine levels, although patients show considerable variability in their response to treatment ^3^. Historically, the modulation of beta-band activity has emerged as a consistent neurophysiological marker of dopaminergic treatment. Recent investigations have moved beyond sustained activity to demonstrate that levodopa alters transient network dynamics; for example, Tinkhauser et al. ^4^ revealed that the OFF state is characterized by prolonged pathological "beta bursts" that are truncated following treatment. Despite its therapeutic relevance, STN beta activity does not fully satisfy the requirements of a reliable individual-level biomarker. Its detectability is inconsistent: in a multicentre study of 156 patients, artifact-validated beta peaks were identified in only 65.59% of hemispheres and bilaterally in 47.25% of patients ^5^. Moreover, beta bursts explain only a limited-to-moderate proportion of symptom variability, with reported values of 0.4–0.5 and estimates as low as 0.18–0.36 in the largest study to date ^6^. As such, the notion that the effects of local alterations might reverberate throughout the brain has led to the use of large-scale recordings. Consistent with a more distributed representation of clinical state, recent artificial-intelligence approaches decoded and predicted symptoms substantially more accurately from electrocorticographic signals than from deep-brain recordings ^7^. Magnetoencephalography (MEG) can provide an optimal non-invasive window into these dynamics. The sub-millisecond temporal resolution of MEG, combined with advanced source reconstruction techniques ^8–108–10^, has enabled the detailed investigation of both localized sensorimotor circuits and distributed whole-brain networks in PD ^11,12^.

For instance, Levodopa-induced increases in sensorimotor cortical beta power have been associated with improvements in akinesia and rigidity ^13^. Furthermore, the restoration of beta oscillations in cortical regions with high dopamine receptor density has been shown to correlate with reduced motor symptom severity, as measured by the Unified Parkinson’s Disease Rating Scale (UPDRS-III) ^14^. Expanding beyond isolated frequency bands, Peña and colleagues ^15^ pointed out that subject-specific broadband spectral profiles are essential for accurately classifying a patient’s medication state (OFF vs. ON). Together, these findings suggest that clinically relevant information is not fully captured by local beta dynamics but depends on the broader state of the brain network. This motivates complementary, network-level approaches that are neither confined to a predefined frequency band nor based exclusively on local activities. High-temporal-resolution approaches have begun to capture this phenomenon, revealing differential medication effects on time-resolved spectral connectivity ^16^ and the spatial spread of neuronal avalanches ^17^. Furthermore, the loss of dopaminergic tone shifts the whole-brain dynamics to a stereotyped regime, characterized by a restricted functional repertoire ^18^.

To characterize, at the individual level, the different whole-brain dynamical regimes induced by medication, we conducted a repeated-measures MEG study utilizing a source-space microstate framework ^19^ in a selected, homogeneous cohort of PD patients. In short, we focused on peaks of the overall brain activity and clustered them into different states. Then, we estimated the probability of transitioning from any two states, thence defining subject-specific transition matrices before (OFF) and after (ON) medication. In this light, we treated the transition probabilities of the unmedicated state as a baseline measure of pathological dynamics and hypothesized that effective dopaminergic intervention would reconfigure the transition probabilities in proportion to clinical efficacy.

## Results

### Participants

Demographic and clinical characteristics of the study cohort are summarized in Table 1. The final analysis included 13 patients with Parkinson’s disease (12 males; age range: 40–81 years; mean education: 10.92 years). Disease duration ranged from 9 to 180 months, encompassing Hoehn and Yahr (H&Y) stages I through IV. Clinical motor evaluations, assessed via the UPDRS-III (excluding tremor sub-items), yielded a mean score of 28.92 (*SD* = 10.61) in the unmedicated OFF state. Approximately 45 minutes following dopaminergic medication intake, the mean UPDRS-III score improved to 18.69 (*SD* = 7.84) in the ON state. Patients were all classified as bradykinetic.

**Table 1.**

|  | **Mean (std)** | **Range min/max** |
| --- | --- | --- |
| **Age** | 65.46 (12.50) | 40 - 81 |
| **Education** | 10.92 (3.32) years | 5 - 17 |
| **Sex** | 12 M / 1 F |  |
| **Disease Duration\*** | 96.75 (55.68) months | 9 - 180 |
| **MMSE** | 28.20 (1.52) | 25.97 - 31-03 |
| **BDI\*** | 7.91 (10.61) | 1 - 38 |
| **H&Y** | 2.23 (0.88) | 1 - 4 |
| **UPDRS-III OFF** | 28.92 (10.61) | 9 - 46 |
| **UPDRS-III ON** | 18.69 (7.84) | 4 - 29 |

\* missing information from one subject

### Microstate computation

To establish a baseline for dynamic network reconfiguration, we first defined subject-specific microstate topographies utilizing a within-subject clustering framework. Each participant underwent four continuous magnetoencephalography (MEG) recordings: two distinct acquisitions in the dopamine-depleted state (OFF) and two following dopaminergic medication (ON). To ensure unbiased intra-subject comparisons, the data were strictly length-matched across the four acquisitions for each participant, yielding a uniform average acquisition duration of 158.02 s (*SD* = 33.79; range: 88.07–197.70 s).

From each acquisition, we extracted the 3,000 peaks with the highest Global Field Power (GFP), defined as those with the highest spatial standard deviation across the 90 anatomically defined regions of interest (ROIs). This yielded a pooled dataset of 12,000 discrete GFP peaks per participant, which was subsequently submitted, separately for each participant, to a modified *k*-means clustering algorithm to empirically derive microstate centroids across k ∈ [3, 20]. Across all participants and k’s, the resulting microstate models captured a robust proportion of the data variance, with Global Explained Variance (GEV) ranging from 0.15 (obtained for a participant at k=3) to 0.54 (obtained for a participant at k=20). The mean GEV across participants and all k’s was 0.34 (*SD* = 0.07).

Following centroid identification, the spatial topographies for each GFP peak of the four acquisitions were competitively assigned to the subject-specific microstate maps based on maximal spatial correlation. This assignment allowed us to compute the empirical transition probability matrices for each condition. Each matrix contains the k states in rows and columns, and the probabilities of transitioning from one state to the next as entries. Finally, to ensure that the observed transition dynamics reflected genuine dynamics rather than static properties, the empirical matrices were z-scored against a null distribution. This distribution was generated using 100 multivariate phase-randomized surrogates per acquisition, preserving both the empirical power spectrum and cross-spectral phase relationships. Hence, the matrices no longer contain the empirical transition probability; they now contain the probability of observing a particular transition probability relative to what is expected under random dynamics (i.e., the phase-preserving surrogates). We observed highly robust medication-induced shifts in the microstate dynamics across all k’s. The remainder of the main text focuses specifically on k=8, as this is the most parsimonious parametrization that captures the dynamical shifts. Comprehensive analyses demonstrating the stability of our findings across the full range of k’s are provided in the supplementary materials (<u>Supp 1</u>).

### Intra-condition and Inter-condition transition matrix similarity

To characterize the stability and reconfiguration of whole-brain dynamics, we analyzed the transition probability matrices across four recording sessions per participant: two acquisitions in the dopamine-depleted (OFF) state and two following administration of dopaminergic therapy (ON). We hypothesized that the transition dynamics in the OFF state would reflect the pathological baseline. Conversely, we expected that changes in the topologies of the transition matrices between the ON and OFF states would index therapy-induced clinical improvement.

Representative transition matrices for a single subject are illustrated in Figure 2. Visual inspection of these matrices reveals high intra-condition consistency in the OFF state, as particularly evident, for example, in the transition probabilities for microstates 6 and 7, and in the ON state.

**Figure 1.**
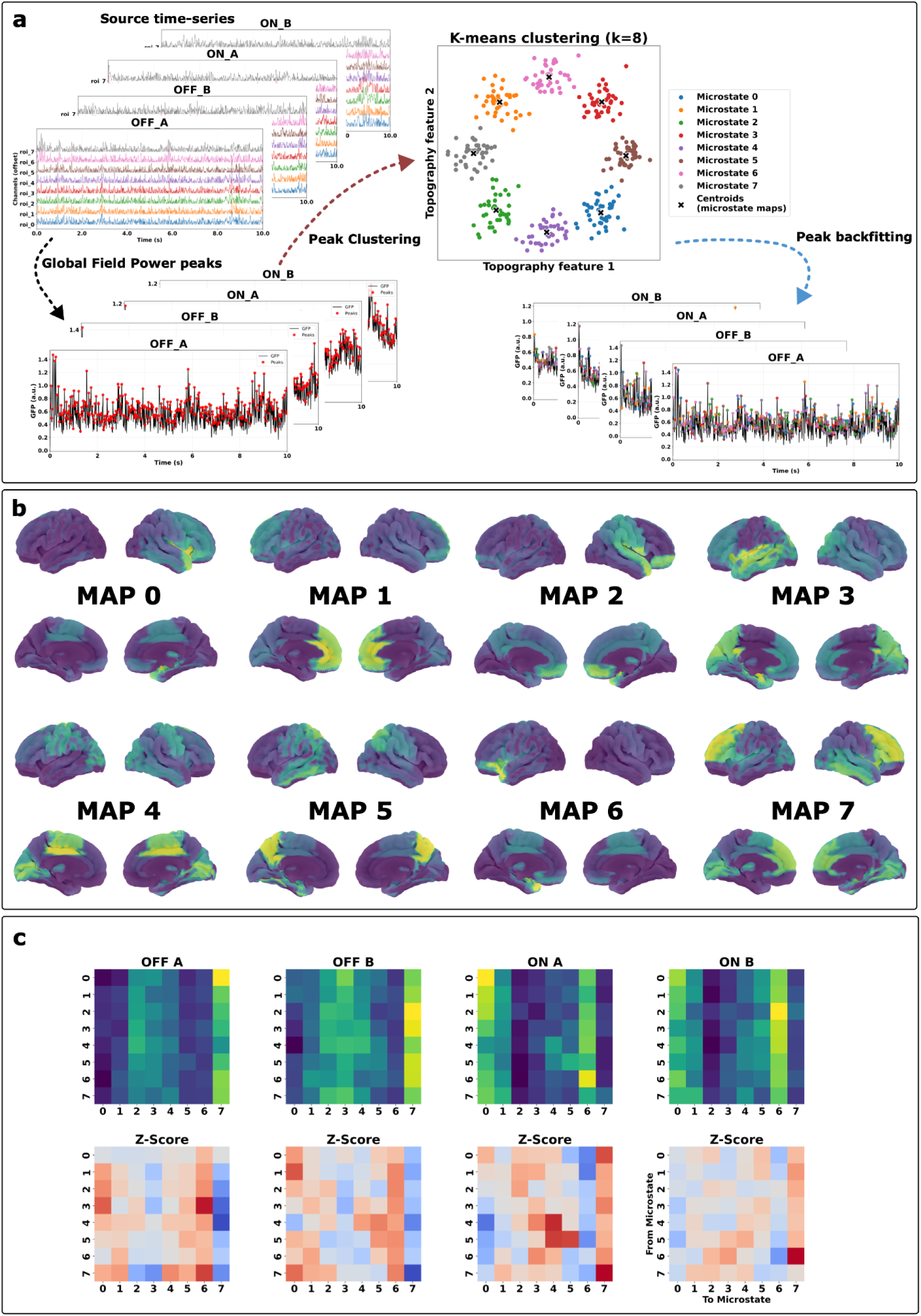
Subject-specific methodological pipeline for microstate transition analysis. (a) Source-reconstructed MEG time series for a representative participant across four recording sessions: two acquisitions (A and B) in the unmedicated (OFF) condition, and two in the medicated (ON) condition. Corresponding Global Field Power (GFP) traces derived from the source-level data for each of the four acquisitions. GFP peaks, pooled across all four acquisitions, were submitted to a modified k-means clustering algorithm to identify a subject-specific set of k=8 prototypical topographies (microstate maps), each represented in a low-dimensional topography feature space with its corresponding centroid. Each GFP peak was then back-fitted to these microstate maps by assigning it to the template with which it showed the highest spatial correlation, yielding a discrete microstate label for each peak per acquisition. (b) Subject-specific microstate topographies (illustrated for k=8). These prototypical states were identified by pooling the GFP peaks across all four acquisitions, ensuring a unified topographical framework for the individual. (c) Derivation of the transition probability matrices. The top row displays the empirical transition matrices mapping the microstate sequences for each acquisition. The bottom row presents the z-scored transition matrices, which have been normalized against a null distribution generated from phase-shuffled surrogate data to isolate non-random transition dynamics.

**Figure 2.**
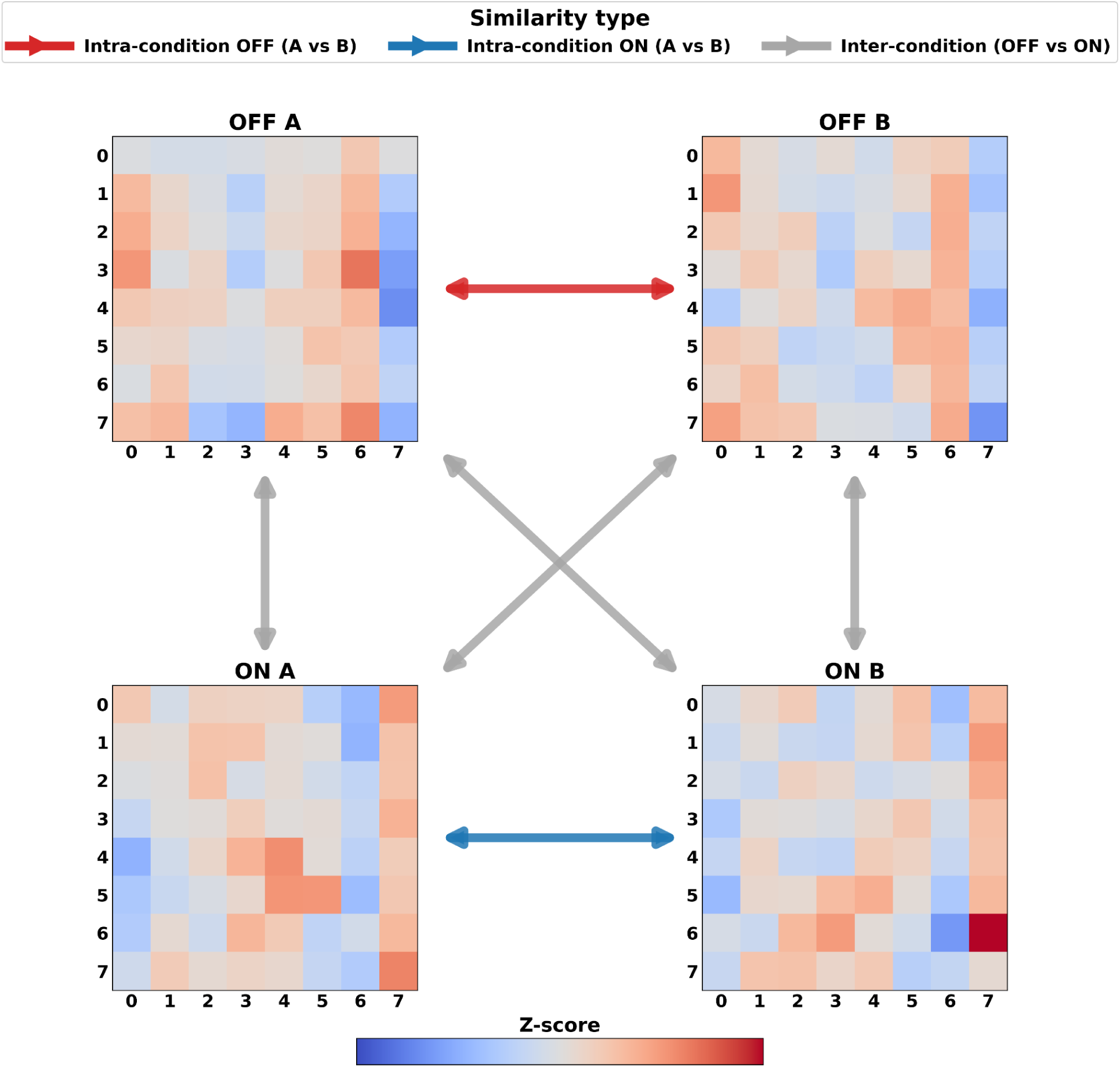
Microstate transition dynamics and topological similarity across pharmacological states. Representative transition probability matrices for a single participant at k=8. The matrices illustrate state-to-state transition sequences (from row to column) across all four recording acquisitions: the dopamine-depleted (OFF) condition (acquisitions A and B) and the medicated (ON) condition (acquisitions A and B). Z-score transition matrices computed against the surrogate transition matrices are shown. Intra-condition topological similarity, quantified via Spearman’s rank correlation coefficient (ρ) between transition matrices of acquisitions within the same pharmacological state (Intra-OFF red and Intra-ON blue). Inter-condition topological similarity, computed as Spearman’s (ρ) across all possible pairwise combinations of unmedicated and medicated acquisitions (gray lines). This panel illustrates the cross-state network reconfiguration induced by dopaminergic therapy.

To quantify this topological similarity, we calculated the Spearman rank correlation coefficient ρ between the z-scored transition matrices of all possible acquisition pairs. Figure 3 displays the resulting similarity distributions across intra-condition (OFF A vs. OFF B; ON A vs. ON B) and inter-condition (OFF vs. ON) pairings.

**Figure 3.**
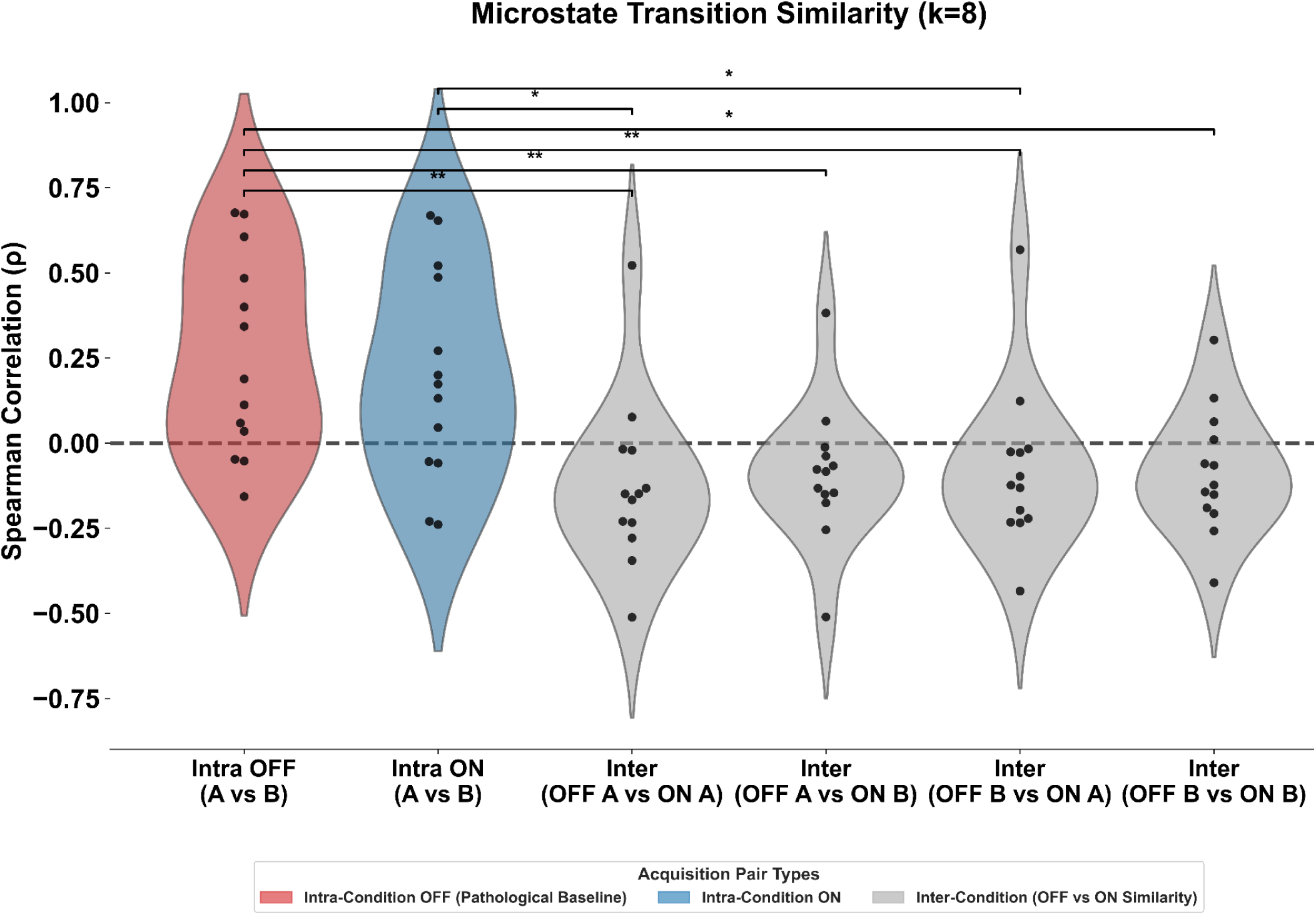
Topological reliability and inter-state similarity of microstate transition matrices. Violin and swarm plots illustrate the distribution of similarity scores across intra-condition and inter-condition acquisition pairs (k=8). Network similarity is quantified using the Spearman rank correlation coefficient ρ between the z-scored transition matrices of different acquisitions. The red distribution represents the highly stable intra-condition baseline in the unmedicated state (Intra OFF), whereas the blue distribution reflects the intra-condition reliability following dopaminergic intervention (Intra ON). Gray distributions denote the cross-condition topological resemblance (Inter-condition). Black horizontal brackets indicate significant differences between distributions, assessed via paired Wilcoxon signed-rank tests. All p-values are corrected for multiple comparisons across the matrix using the Benjamini-Hochberg False Discovery Rate (FDR) procedure (* p < 0.05; p < 0.01; *** p < 0.001). The bold dashed line indicates zero correlation.

The analysis revealed that the intra-condition similarity distributions exhibited the highest correlation coefficients, indicating that the brain steadily dwells in two distinct dynamical regimes as a function of dopamine levels. Conversely, the significantly lower correlations across all inter-condition combinations indicate a change in dynamical regime occurring between the medicated and unmedicated states. Collectively, these findings suggest that the brain appears locked into a (presumably) deleterious dynamics during the OFF-state, and that dopaminergic therapy causes a reconfiguration of state-to-state transition probabilities.

Finally, the identified dynamical regimes were stable over a wide range of k values (*k* = 8–20). Comprehensive statistical reports and transition matrix visualizations for all remaining k parameters are provided in the supplementary materials (<u>Supp 2</u>).

### Clinical Significance of Transition Matrix Reconfiguration

Having demonstrated a systematic reconfiguration of microstate transition probabilities between the OFF and ON states, we next investigated the clinical relevance of this topological shift. We defined a SHIFT_OFF metric, representing the magnitude of brain dynamic reconfiguration, as the difference between the intra-condition similarity between the two OFF-state acquisitions and the average inter-condition similarity (shown in the gray violin plots in Figure 3). We then correlated the SHIFT_OFF with the percentage improvement in motor performance, as assessed by the Unified Parkinson’s Disease Rating Scale (UPDRS) motor score (calculated excluding tremor-related sub-items).

Figure 4 illustrates the relationship between therapy-induced topological reconfiguration and therapeutic responsiveness. The Spearman rank correlations are reported across the entire explored range of k’s (k=3–20). We observed a robust, positive correlation between the degree of medication-induced transition reconfiguration and the magnitude of clinical motor improvement. The inset shows the scatterplot of the correlation corresponding to k=8. The scatterplots for the other k’s are reported in <u>Supp 2</u>. These findings suggest that patients who exhibit a more pronounced departure from the dynamics observed in the dopamine-depleted state gain greater clinical benefit from dopaminergic therapy, highlighting the link between network reconfiguration and therapeutic efficacy in Parkinson’s disease.

**Figure 4.**
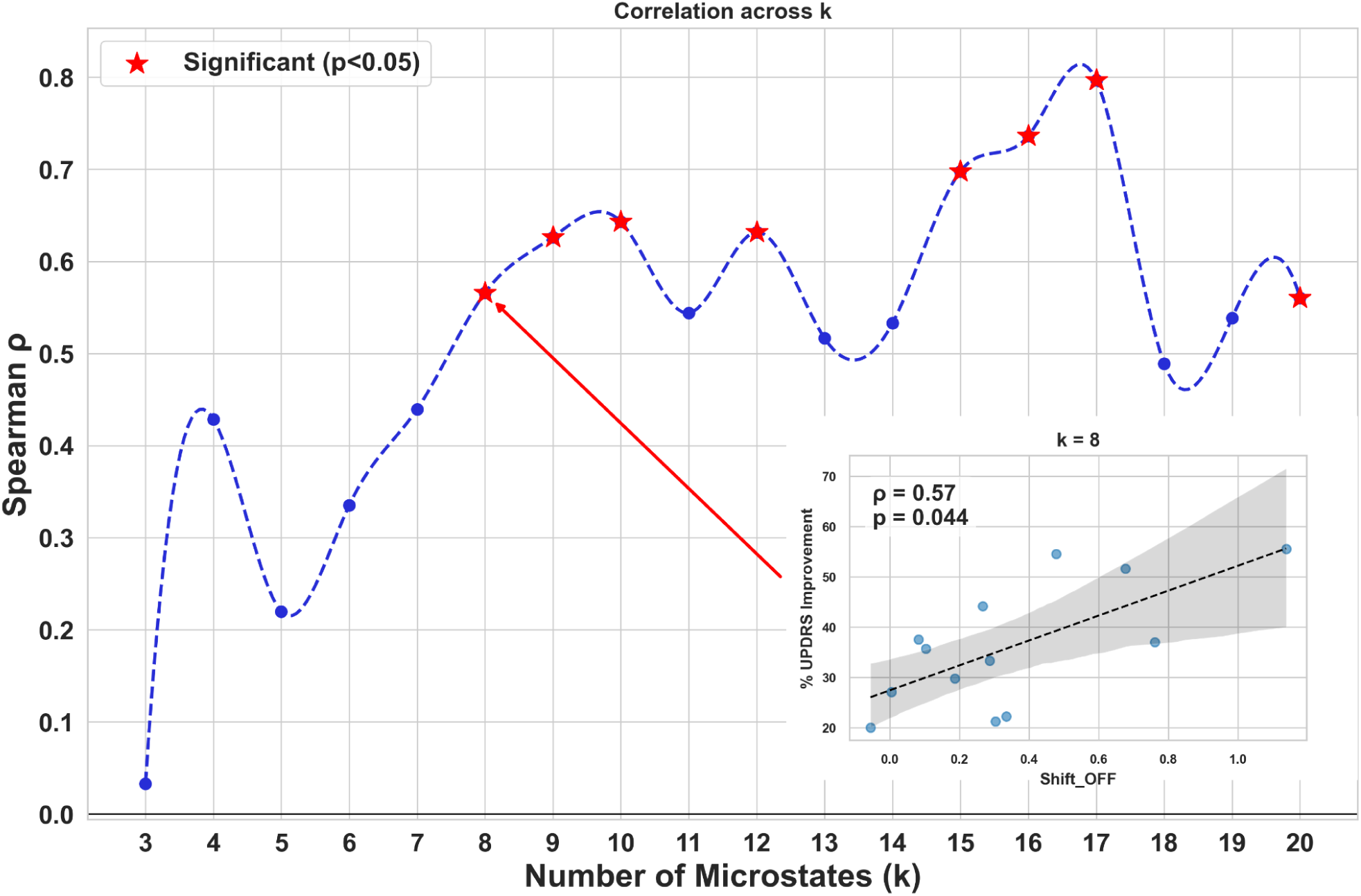
Clinical significance of the pathological transition shift across k’s. The main plot summarizes the Spearman rank correlation ρ across all tested values of k = [3, 20]. Red stars indicate significant correlations (p < 0.05). The inset displays the correlation between the SHIFT_OFF metric—defined as the difference between intra-OFF similarity and the average inter-condition similarity—and the percentage of clinical improvement in UPDRS scores following dopaminergic treatment, for k=8, as a scatter plot.

## Discussion

In the present study, we investigated the impact of dopaminergic therapy on the macroscale brain dynamics of patients with Parkinson’s disease (PD). Using subject-specific microstate transition probabilities derived from source-reconstructed MEG recordings, we identified dynamical shifts across pharmacological states. Specifically, we observed that the transition repertoires indexed the dopamine levels (OFF vs. ON states). Brain dynamics at the individual level were consistent within the ON and OFF states, whereas they differed markedly when comparing the two states directly. The magnitude of the difference between the intra-condition similarity in the OFF state minus the average inter-condition similarities was proportional to the clinical motor improvement (UPDRS). Together, these findings suggest that therapeutic responsiveness in PD is closely linked to a shift in the structure of the whole-brain dynamics.

We observed high intra-condition topological reliability within both the unmedicated (OFF) and medicated (ON) states. However, significant clinical correlations were obtained exclusively with the magnitude of network reconfiguration—specifically, the divergence of the inter-condition topology from the stable intra-OFF-state baseline. These results imply that therapeutic responsiveness is not strictly related to a “universally optimal” functional state. Instead, the magnitude of clinical improvement following drug administration appears to be most strongly associated with the extent to which treatment shifts neural activity away from the dynamical regime characteristic of the OFF-medication state.

In this study, we moved beyond conventional analyses of periodic neural activity ^11,12^ by investigating brain dynamics without prespecifying a frequency band of interest. Support for this approach comes from both experimental and modeling studies. Experimentally, levodopa-induced neurophysiological changes have been captured at the individual level through alterations in the propagation of neuronal avalanches, which exhibit scale-free properties ^17^. Complementarily, modeling work has shown that changes in the spatial distribution of aperiodic activity observed in patients can be successfully reproduced as a function of levodopa levels ^20^.

The microstate framework ^21^ was selected to investigate the spontaneous transient dynamics of whole-brain neuronal networks in PD ^22,23^. Previous studies utilizing classical electroencephalographic (EEG) microstates and their properties (e.g., occurrence, time coverage, duration) have successfully differentiated PD patients from healthy controls ^24–26^. Moreover, microstate sequences have proven superior to simple sliding-window approaches for classifying PD patients and controls based on brain network dynamics, effectively serving as a temporal scaffold to guide the computation of time-averaged functional connectivity ^27^. Spectrally resolved microstate analyses have also successfully identified cognitive phenotypes within PD, distinguishing between patients with and without mild cognitive impairment ^28^.

Methodologically, the present study builds upon the source-level MEG microstate pipeline proposed by Tait and Zhang (2022). However, our approach introduces two critical modifications. First, to capture the highly individualized nature of dopaminergic responsiveness ^15^, we computed subject-specific microstate topographies rather than relying on a group-level grand average. This circumvents the well-documented heterogeneity of "canonical" EEG microstates across cohorts ^29^ and avoids the methodological ambiguities associated with template normalization ^30^. Second, transition matrices were derived strictly from Global Field Power (GFP) peaks, bypassing the traditional back-fitting procedure to the continuous recording. This conservative, data-driven choice was made to simplify the analytical pipeline and mitigate the lack of consensus regarding optimal back-fitting strategies ^31^.

Despite these methodological simplifications, GFP-derived transition matrices proved highly sensitive to dopaminergic modulation. This aligns with prior EEG evidence demonstrating that dopamine therapy alters microstate dynamics, syntax and transition probabilities ^32,33^. The consistency of our findings across a wide range of k’s further underscores the reliability of the proposed framework. Importantly, transition probability matrices have not only successfully differentiated the dynamics of healthy controls from those of PD patients (Guo et al. 2024) but also closely track clinical severity scores ^34,35^.

This work has limitations. First, the sample size is relatively small. However, this was a deliberate consequence of restricting our cohort to the bradykinetic PD subtype to minimize the confounding influence of phenotypic heterogeneity on dopaminergic responsiveness. Second, variations in disease duration and levodopa-equivalent daily dose were not explicitly incorporated into the model and may contribute to unexplained variance. Third, regarding experimental design, the intra-OFF acquisitions (A and B) were collected closer in time than the inter-condition acquisitions (OFF vs. ON), raising the potential confound of temporal proximity. However, the strong correlation with clinical improvement makes it unlikely that our results are solely a spurious result due to the different time intervals. Finally, cognitive and physical fatigue over the course of the recording sessions ^36^ may have subtly influenced the transition dynamics, thereby representing a latent variable that is difficult to disentangle.

In conclusion, this study offers two core insights. First, the unmedicated bradykinetic brain operates within a rigid, stereotyped dynamical attractor. Second, quantifying the topological shift in macro-scale state transitions away from this pathological baseline provides a robust, mechanistically grounded biomarker for evaluating personalized pharmacological responses in Parkinson’s disease.

## Methods

### Participants and Clinical Characterization

#### Recruitment and Diagnostic Criteria

A cohort of consecutive patients with early-stage Parkinson’s Disease (PD) was recruited from the Movement Disorders Unit of the First Division of Neurology at the University of Campania “Luigi Vanvitelli” (Naples, Italy). Diagnostic status was confirmed in accordance with the UK Parkinson’s Disease Society Brain Bank clinical diagnostic criteria ^37^. To ensure cohort homogeneity and minimize confounding variables, the following inclusion criteria were applied: age of onset > 40 years, specifically to exclude early-onset parkinsonism genotypes.

#### Exclusion criteria

Participants were excluded based on: a diagnosis of PD-associated dementia according to established consensus criteria ^38^, mini-mental state examination (MMSE) < 24 and the presence of comorbid neurological disorders or any unstable systemic medical conditions that could interfere with the study’s primary outcomes.

#### Clinical and Pharmacological Assessment

Disease severity and motor disability were quantified using the UPDRS Part III ^39^ (Motor Examination) and H&Y staging ^40^. To capture the effect of therapy, motor assessments were performed in the ‘off-state’ (a night of therapy withdrawal) and in the ‘on-state’ at least 45 minutes after medication intake.

#### Ethical Considerations

All participants provided written informed consent prior to inclusion. The study protocol was formally approved by the Institutional Review Board (Local Ethics Committee) of the University of Campania “Luigi Vanvitelli” and was conducted in strict adherence to the ethical principles of the Declaration of Helsinki.

#### Meg recordings

Magnetoencephalographic (MEG) recordings were acquired using a 163-magnetometer system (AtB Biomag UG, Ulm, Germany) with nine reference sensors, housed within a magnetically shielded room. Prior to acquisition, four head-position coils and four anatomical fiducials (nasion, bilateral pre-auricular points, and apex) were digitized via a Polhemus Fastrak system. Resting-state data were collected over four acquisitions, each lasting at least 3.5 minutes, with two acquisitions while participants were in a medication-off state and two in a medication-on state. Participants without the two acquisitions per condition were excluded. Subjects were instructed via intercom to remain relaxed with eyes closed and avoid specific mental activity; head position was verified at the onset of each recording block. Data were digitized at a sampling frequency of 1024 Hz after applying an analog anti-aliasing filter. Offline preprocessing was performed in MATLAB using the FieldTrip toolbox (v.2014)^41^, where signals were constrained to a 0.5–48 Hz bandwidth using a 4th-order Butterworth IIR band-pass filter. Electrocardiogram (ECG) and electrooculogram (EOG) data were recorded concurrently to facilitate subsequent artifact identification.

#### MRI acquisition

Structural MRI was conducted on a 3-T system (General Electric Healthcare, Milwaukee, WI, USA) using an 8-channel parallel head coil. Imaging was performed either post-MEG or within a 21- to 30-day window preceding the session. Anatomical characterization used a 3D T1-weighted inversion-recovery-prepared fast spoiled gradient-recalled echo (IR-FSPGR) sequence with the following parameters: TR = 6988 ms, TI = 1100 ms, TE = 3.9 ms, flip angle = 10°, and voxel size = 1 × 1 × 1.2 mm³.

#### Preprocessing

Environmental noise was attenuated via Principal Component Analysis (PCA) by orthogonalizing reference signals and subtracting their projections from the MEG sensors ^42^ using the FieldTrip toolbox implementation ^41^. Following expert visual inspection to exclude noisy data segments, an average of 130 ± 2 channels was retained per participant. Finally, Independent Component Analysis (ICA) ^43^ was employed to identify and remove physiological artifacts, specifically targeting cardiac (typically 1–2 components) and ocular (0–1 components) contributions to the MEG signal.

#### Source reconstruction

Source-level activity was reconstructed using the Linearly Constrained Minimum Variance (LCMV) algorithm implemented in the FieldTrip toolbox ^41^. Following fiducial-based co-registration of MEG data to native MRIs, a single-shell volume conduction model ^44^ was employed to estimate broadband time series for the centroids of 116 Automated Anatomical Labeling (AAL) regions ^45,46^. We specifically analyzed the first 90 ROIs, excluding the cerebellum due to lower reconstruction reliability in that region. Finally, all source-space data were visually inspected, yielding high-quality segments for each acquisition to ensure the absence of residual artifacts. Segments shorter than 2 seconds were excluded.

#### Microstate computation

The analytical pipeline was adapted from the source-space microstate framework of Tait et al. (2022) ^19^, utilizing LCMV reconstruction in conjunction with the AAL-90 atlas. Source time-series were band-pass filtered (2–30 Hz) and standardized via z-scoring. Global Field Power (GFP)—calculated as the spatial standard deviation across ROIs—was smoothed using a five-sample moving average to ensure robust peak detection. To account for potential source-orientation flipping, absolute values of the z-scored estimates were extracted at each GFP peak. For each condition (OFF and ON medication), we had two acquisitions (A and B). We cropped each acquisition to its minimum length, separately for each participant. Then, we extracted the 3000 highest GFP peaks for each acquisition from each participant (12000 GFP per participant), enforcing a minimum inter-peak interval of 10 samples (∼0.01s).

The resulting aggregate of GFP peaks was submitted separately per participant to a modified k-means clustering algorithm using the Pycrostates Python package ^47^. The clustering procedure involved 50 random initializations, with the solution maximizing Global Explained Variance (GEV) selected for further analysis. Final cluster centroids were refined through eigenvector decomposition. We explored the number of microstates in the range k ∈[3, 20].

For each k, after the subject-specific microstate maps were computed, GFP peaks for all four acquisitions (OFF_A, OFF_B, ON_A, and ON_B) were competitively assigned to the identified centroids based on spatial correlation and used to compute the transition matrices (i.e., transition matrices based only on the GFP peaks) (see Figure 1). For each acquisition, a row-normalized transition probability matrix was computed, encompassing both cross-state and self-state transitions.

To ensure that the observed transition dynamics reflected genuine non-random state sequencing rather than underlying spectral autocorrelations, the empirical transition matrices were standardized against a null distribution. Specifically, we generated phase-shuffled surrogates for each acquisition (100 surrogates for each acquisition). By applying a phase-randomization procedure that independently shuffles the original data while preserving both the power spectrum and cross-spectral phase relationships. Each empirical transition probability was subsequently z-scored against this surrogate distribution, yielding a normalized transition matrix for each acquisition.

To quantify the topological similarity of the transition matrix, we computed the Spearman rank correlation coefficient of z-scored transition matrices across different acquisitions. Specifically, we computed the following metrics:

● **Intra-condition similarity**: Calculated as the correlation between the two acquisitions within the same pharmacological state (Intra OFF: OFF A vs. OFF B; and Intra ON: ON A vs. ON B). We hypothesize that a highly consistent Intra_OFF matrix may reflect the pathological baseline of the dopamine-depleted brain.
● **Inter-condition similarity**: Calculated as one of the four possible cross-condition cross-correlations (OFF A vs. ON A, OFF A vs. ON B, OFF B vs. ON A, and OFF B vs.

ON B). This metric captures the topological shift between the unmedicated and medicated states.

#### Statistical analysis

To statistically evaluate the condition-dependent shifts for the transition matrices, we compared the derived similarity distributions (*Intra OFF*, *Intra ON*, and *Inter OFF/ON*) at the group level with non-parametric statistical methods.

Differences in topological patterns between conditions were assessed using paired, two-sided Wilcoxon signed-rank tests. This allowed for direct within-subject comparisons across the three primary metric pairings (e.g., *Intra OFF* vs. *Intra ON*; *Intra OFF* vs. *Inter OFF/ON*; and *Intra ON* vs. *Inter OFF/ON*).

To control the Type I error rate across all pairwise comparisons, for each k, all resulting *p*-values were corrected for multiple comparisons using the Benjamini-Hochberg False Discovery Rate (FDR) procedure. A corrected alpha level of *p* < 0.05 was established as the threshold for statistical significance.

Finally, to test our primary clinical hypothesis, we utilized the Spearman rank-order correlation to assess the relationship between the magnitude of transition matrix reconfiguration (calculated as the scalar difference between the baseline *Intra OFF* and the average inter-condition *Inter_OFF/ON*) and the patient’s relative clinical benefit, defined as the percentage improvement (OFF vs ON) in the Unified Parkinson’s Disease Rating Scale (UPDRS) motor score. Motor scores were computed excluding the items related to tremor assessment.

## Data availability

The MEG source microstate data are available upon reasonable request to the corresponding author, conditional on appropriate ethics approval at the local site.

## Code availability

All the code for the analysis is available at https://github.com/suforraxi/microstate_pd_onoff.git

## Acknowledgments

This work was funded by:

Governo Italiano Ministero per lo sviluppo Economico, ACCORDI PER INNOVAZIONE. Approccio User-friendly integrato per Diagnosi, Assistenza e Cura Efficaci—AUDACE. CUP: B69J23006050007.

European Union “NextGenerationEU”, (Investimento 3.1.M4. C2), project IR0000011, EBRAINS-Italy of PNRR

## Author contributions

M.D., G.S., P.S. conceived the experiment; M.D., M.A., P.S. designed the methodology; M.D. and M.A. analyzed the data; M.D. wrote the first draft; M.D., M.A., E.T.L., M. De Luca, E.G., D.T., C.G., D.D, G.S. and P.S. revised and wrote the final manuscript.

## Disclosure of Delegation to Generative AI

The authors declare the use of generative AI in the research and writing process. According to the GAIDeT taxonomy (2025), the following tasks were delegated to GAI tools under full human supervision:

- Code generation
- Code optimization
- Summarizing text
- Adapting and adjusting emotional tone
- Reformatting

The GAI tool used was: ChatGPT4o, Gemini 3, Claude Sonnet 5.

Responsibility for the final manuscript lies entirely with the authors.

GAI tools are not listed as authors and do not bear responsibility for the final outcomes.

Declaration submitted by: ’Collective responsibility’

