## Supplementary for "Departure from OFF-State Microstate Dynamics Tracks Levodopa Response in Parkinson’s Disease"

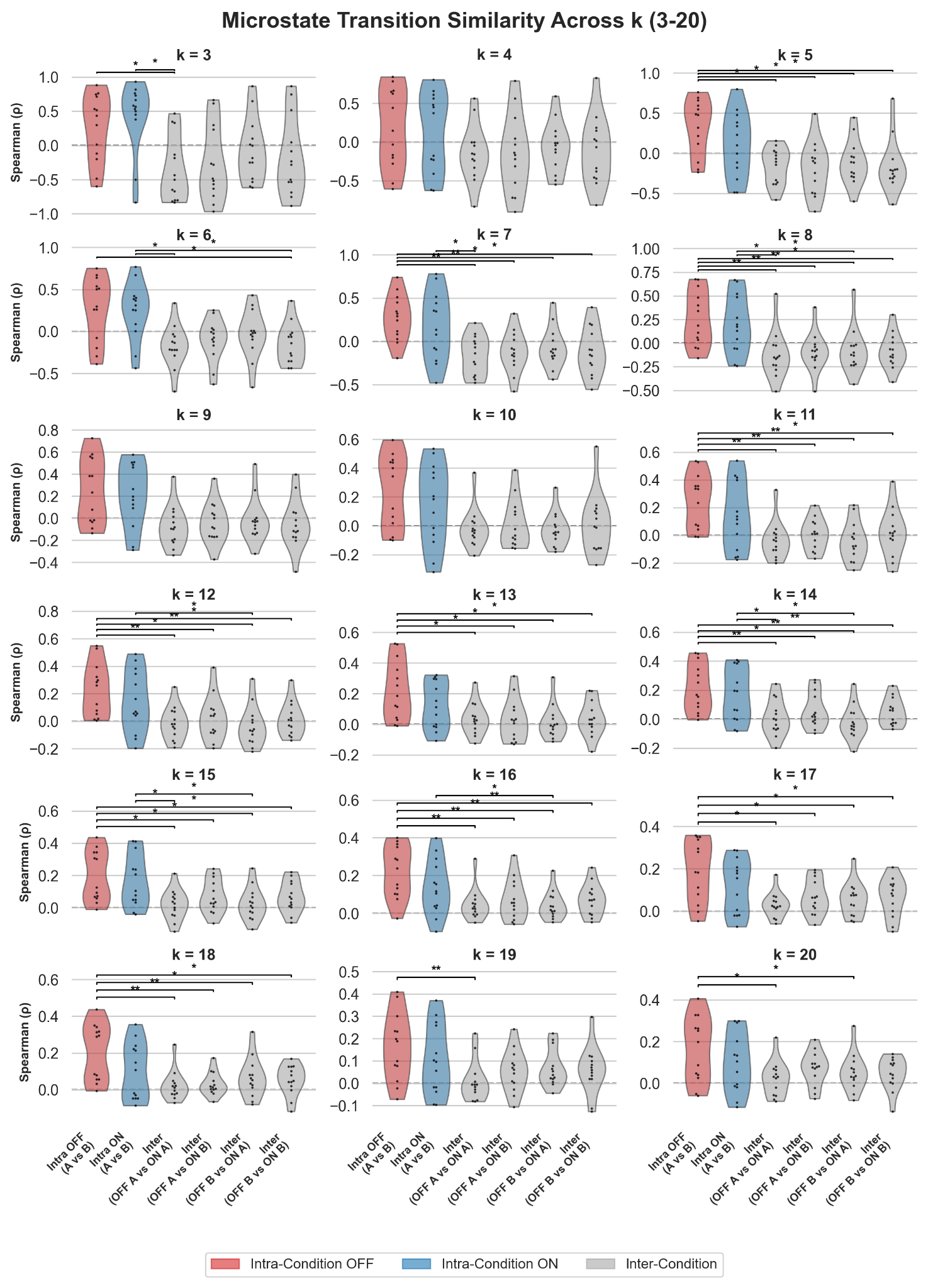

#### ***Supplementary Figure 1. Topological intra-condition similarity and inter-condition similarity of microstate transition matrices across k ∈ 3 to 20. Violin and swarm plots illustrate the distribution of similarity scores across intra-condition and inter-condition acquisition pairs for each model order (k). Network similarity is quantified using the Spearman rank correlation coefficient ρ between the z-scored transition matrices of different acquisitions. Within each k, the red distribution represents the highly stable intra-condition baseline in the unmedicated state (Intra OFF), whereas the blue distribution reflects the intra-condition reliability following dopaminergic intervention (Intra ON). Gray distributions denote the cross-condition topological resemblance (Inter-condition). Black horizontal brackets indicate significant differences between distributions at a given scale, assessed via paired Wilcoxon signed-rank tests. All p-values are corrected for multiple comparisons using the Benjamini-Hochberg False Discovery Rate (FDR) procedure (*p < 0.05; **p < 0.01; *** p < 0.001). A bold dashed line indicates zero correlation.***

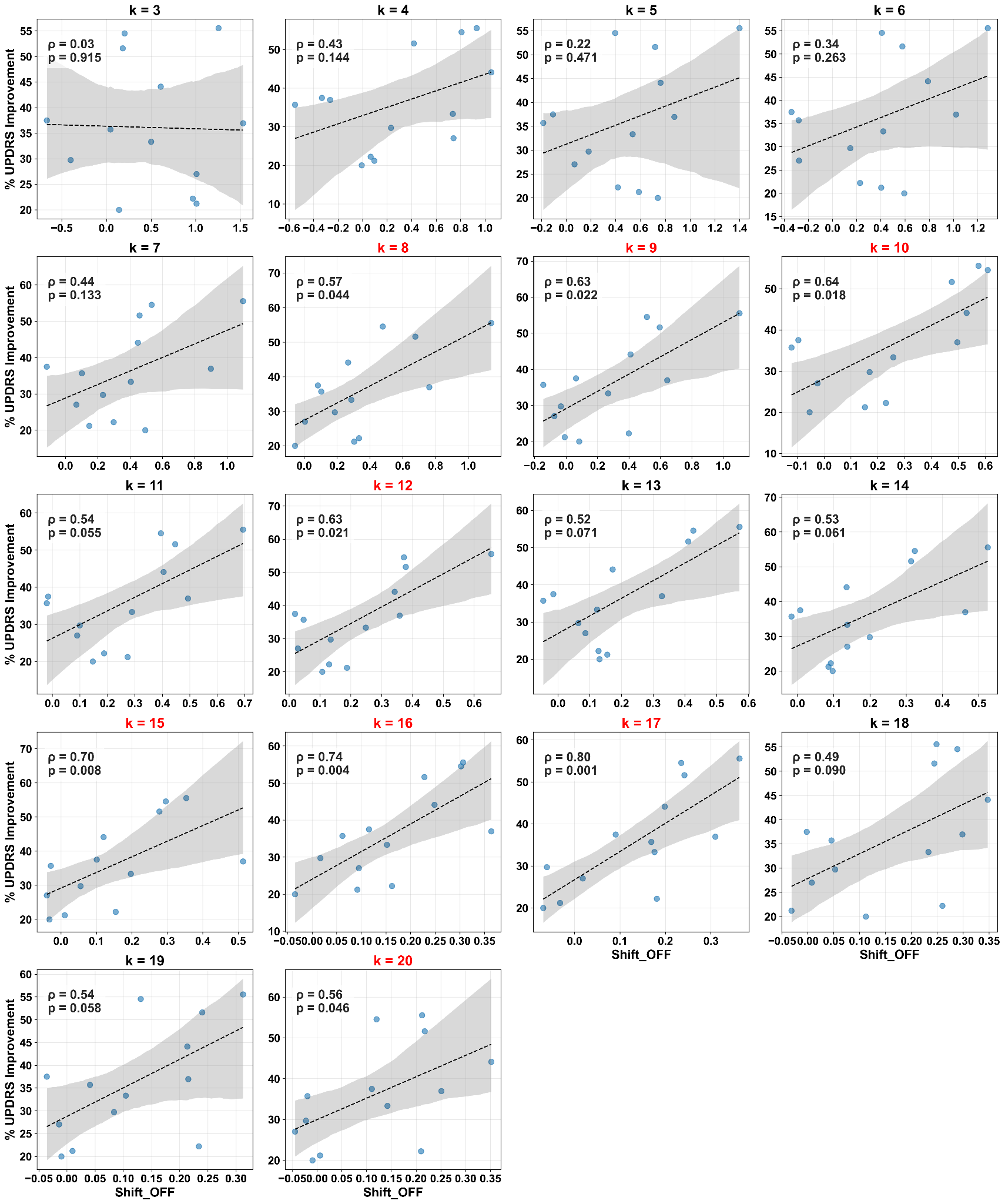

#### ***Supplementary Figure 2: Clinical significance of the pathological transition shift across different k parameters for the k-means algorithm. Scatter plots detail the relationship between the SHIFT_OFF metric—defined as the difference between the intra-condition OFF stability and the average inter-condition similarity—and the percentage of clinical improvement in UPDRS motor scores following dopaminergic treatment. This correlation is illustrated comprehensively for all k ∈ [3, 20]. Panels demonstrating a statistically significant correlation (p < 0.05) are highlighted with red titles and regression lines, confirming the robustness of the clinical-topological relationship across the parameter space.***
